# 5-MeO-DMT induces sleep-like LFP spectral signatures in the hippocampus and prefrontal cortex of awake rats

**DOI:** 10.1101/2023.06.01.543303

**Authors:** Annie da Costa Souza, Bryan da Costa Souza, Arthur França, Marzieh Moradi, Nicholy da Costa Souza, Katarina Leão, Adriano Bretanha Lopes Tort, Richardson Leão, Vítor Lopes-dos-Santos, Sidarta Ribeiro

## Abstract

5-methoxy-N,N-dimethyltryptamine (5-MeO-DMT) is a potent classical psychedelic known to induce changes in locomotion, behaviour, and sleep in rodents. However, there is limited knowledge regarding its acute neurophysiological effects. Local field potentials (LFPs) are commonly used as a proxy for neural activity, but previous studies investigating psychedelics have been hindered by confounding effects of behavioural changes and anaesthesia, which alter these signals. To address this gap, we investigated acute LFP changes in the hippocampus (HP) and medial prefrontal cortex (mPFC) of freely behaving rats, following 5-MeO-DMT administration. 5-MeO-DMT led to an increase of delta power and a decrease of theta power in the HP LFPs, which could not be accounted for by changes in locomotion. Furthermore, we observed a dose-dependent reduction in slow (20-50 Hz) and mid (50-100Hz) gamma power, as well as in theta phase modulation, even after controlling for the effects of speed and theta power. State map analysis of the spectral profile of awake behaviour induced by 5-MeO-DMT revealed similarities to electrophysiological states observed during slow-wave sleep (SWS) and rapid-eye-movement (REM) sleep. Our findings suggest that the psychoactive effects of classical psychedelics are associated with the integration of waking behaviours with sleep-like spectral patterns in LFPs.

## INTRODUCTION

Classical psychedelics such as 5-methoxy-N,N-dimethyltryptamine (5-MeO-DMT) comprise structural analogues of serotonin that in humans induce major changes in perception, movement, emotion and cognition (Barsuglia et al., 2018; Davis et al., 2018; Ingvar and Soderberg, 1956; Ornelas et al., 2022; Ott, 2001; Perlin, 1956; Rinkel et al., 1952; Wießner et al., 2023, 2022c, 2022b, 2022a). In rodents, the acute behavioural effects of classical psychedelics include changes in locomotion, space occupancy, and stereotyped behaviours such as wet dog shaking, head twitching, tremors, and backward gaiting (Halberstadt et al., 2008; Kyzar et al., 2016). Rats under acute effects of d-LSD, another serotonin analogue, decreased the number of arms entries in the Y-maze task (Drew et al., 1973), and presented inhibition of fighting in a shock-elicited fighting paradigm when treated with high doses of d-LSD or 5-MeO-DMT (Walters et al., 1978). Head-twitching increases in a dose-dependent manner, while mating ultrasonic vocalizations are suppressed by 5-MeO-DMT (Jefferson et al., 2023). The startle response is altered upon psychedelic dosing in rodents (Davis and Sheard, 1974). Furthermore, a conditioned avoidance response was disrupted progressively over time after NN-DMT dosing, with a peak effect at 8 min post-dosing (Stoff et al., 1977).

More recently, the electrophysiological dynamics that underlie acute psychedelic experiences have begun to be investigated in rodents. The spike rates of mPFC neurons recorded from anaesthetised rats have been shown to increase after dosing of the classical psychedelic DOI (Puig et al., 2003), but another study found that mean firing rates either increased or decreased in different mPFC neurons after DOI dosing, while paired cell synchrony did not change and LFP signals showed decreased power in the gamma frequency band (Wood et al., 2012). DOI also led to a decrease in prefrontal cortical slow oscillations in anaesthetised animals, possibly decreasing global synchronisation (Celada et al., 2008). Likewise, the intravenous dosing of 5-MeO-DMT to anaesthetised rats led to either decreased or increased firing rates in cortical pyramidal neurons, with a corresponding reduction in the power of slow oscillations recorded from the mPFC. Another study showed that 5-MeO-DMT alters the cortico-thalamic activity in freely behaving mice, leading to increased delta power in the V1 cortex, increased beta power in the medial mPFC, decreased beta power in the medial thalamus, and increased gamma power in the medial mPFC, as well as increased coherence in the beta frequency band across the V1 cortex, the medial thalamus and the medial mPFC; and increased coherence in the theta frequency band between the medial mPFC and the medial thalamus (Riga et al., 2018).

Interpreting the electrophysiological findings is complicated by the behavioural confounders such as sleep, anaesthesia, and speed, and by the neuroanatomical constraint of exclusively evaluating cortical signals. Classical psychedelics, like psilocin, have been shown to impact sleep maintenance and cortical oscillations in mice (Thomas et al., 2022), while the mental states they induce in humans share similarities with dreaming (Kraehenmann, 2017; Preller and Vollenweider, 2018; Sanz et al., 2018; Zamberlan et al., 2018). This intriguing paradox remains unresolved. Most relevant studies were conducted on anesthetised animals, which prevents the manifestation of both waking and physiological sleep. In freely moving mice, psilocin delayed REM sleep onset and decreased NREM sleep maintenance, resulting in heightened 4 Hz oscillations (Thomas et al., 2022). An initial report found that 5-MeO-DMT reduced power in the theta frequency range (5-10 Hz) and increased slow wave activity while animals remained awake, suggesting a hybrid state with features of both waking and slow-wave sleep (Bréant et al., 2022). However, this notion is limited by its reliance on cortical data uncontrolled for speed. To date, no electrophysiological studies of psychedelics have controlled for speed or investigated beyond the cerebral cortex (Bréant et al., 2022; Celada et al., 2008; Riga et al., 2018; Thomas et al., 2022).

The hippocampus (HP) plays a key role in rodent navigation and cognition, and its neural activity is characterised by various rhythms. In particular, theta oscillations dominate hippocampal activity during active behaviour and REM sleep (Winson, 1974). Further, theta amplitude is positively correlated with animal speed during waking periods (Ledberg and Robbe, 2011; Whishaw and Vanderwolf, 1973) and strongly influences cortical signals (Sirota et al., 2008). The theta cycle is thought to organise multiple operations in the hippocampal circuit, such as the activation of different pathways (Buzsáki and Moser, 2013; Hasselmo et al., 2002). Additionally, theta-nested gamma oscillations are thought to reflect different circuits within the hippocampal formation (Colgin et al., 2009; Fernandez-Ruiz et al., 2023; Lasztóczi and Klausberger, 2014). Thus, spectral analyses have the potential of capturing acute physiological changes related to the psychedelic experience.

In the absence of data from both the HP and cortex, and without controlling for speed and behavioural variability, it is not possible to draw any conclusions from rodents about the sleep-waking spectral features of LFP signals during acute psychedelic states. To fill this gap, we combined chronic simultaneous LFP recordings from the HP and mPFC with a detailed quantitative analysis of speed as well as stereotyped behaviours after dosing of 5-MeO-DMT. Based on the current literature, we hypothesised that the main brain oscillations should all be altered during the acute psychedelic state. Also, we hypothesised that 5-MeO-DMT would induce an atypical waking state that may spectrally resemble sleep states.

## METHODS

### Animals

A total of 17 adult male rats were used (*Rattus norvegicus*, Wistar, 250-350g, ∼2 months old). The animals were housed in an appropriate vivarium under controlled temperature and humidity, with lights on at 6:00 and lights off at 18:00. This research was previously submitted and approved by the local Ethics Committee CEUA at UFRN (permit #11/2015) and by the Brazilian Health Regulatory Agency (ANVISA AEP # 018/2021).

### Drug

The study used 5-MeO-DMT (Merck) dissolved in ultrapure water (18.2 MΩ.cm).

### Electrode manufacturing

Electrodes were designed and handmade as described elsewhere (França et al., 2020; Souza et al., 2018). The electrode arrays were designed to be implanted in the mPFC (16 channels) and the hippocampus (16 channels) according to Paxinos Atlas (Paxinos, 2013). In each area, two rows of 8 electrodes were implanted along the anteroposterior axis, with 250 mm of inter-electrode space.

### Surgery and post-operative animal care

Experimental animals were subjected to surgery for electrode implantation as described in Souza et al., 2018. Briefly, anaesthesia was induced with inhaled isoflurane at 5% followed by intraperitoneal ketamine and xylazine at respective doses of 100 mg/kg and 8 mg/kg. When needed supplemental doses of ketamine and xylazine were applied to maintain anaesthesia. Electrodes and the cannula were then implanted aiming at the dorsal hippocampus (Anteroposterior (AP): −3.28mm; Mediolateral (ML): 2.0mm; Dorsoventral (DV): 2.2mm); the prefrontal cortex (AP: +3.72mm; ML: −0.5mm; and DV: 3.5mm); and the third ventricle (AP: −0.96mm; ML: 1.8mm; and DV: 3.2mm). Following the surgery animals were treated with analgesic, anti-inflammatory, and antibiotic drugs and given time to recover.

### Experimental design

Animals were submitted to four, weekly-spaced experimental sessions (**Figure 1A**; 1 to 8 sessions per rat, totalling 54 drug and 20 saline sessions). Each experimental session consisted of three consecutive days that started with a one-hour recording of the animal in the arena (55.5 x 65 x 45 cm). On the second day, after the baseline recording, animals were injected with one of the three 5-MeO-DMT doses (ICV groups: 100ug/ul, 150ug/ul, or 200ug/ul; or IP groups: 1mg, 3mg, or 10mg) or saline (referred to as 0g ICV or IP) and recorded for an additional two to three hours. Electrophysiology and animal behaviour were recorded at each session. All experiments reported here refer to the second day of each experimental week.

**Figure 1:**
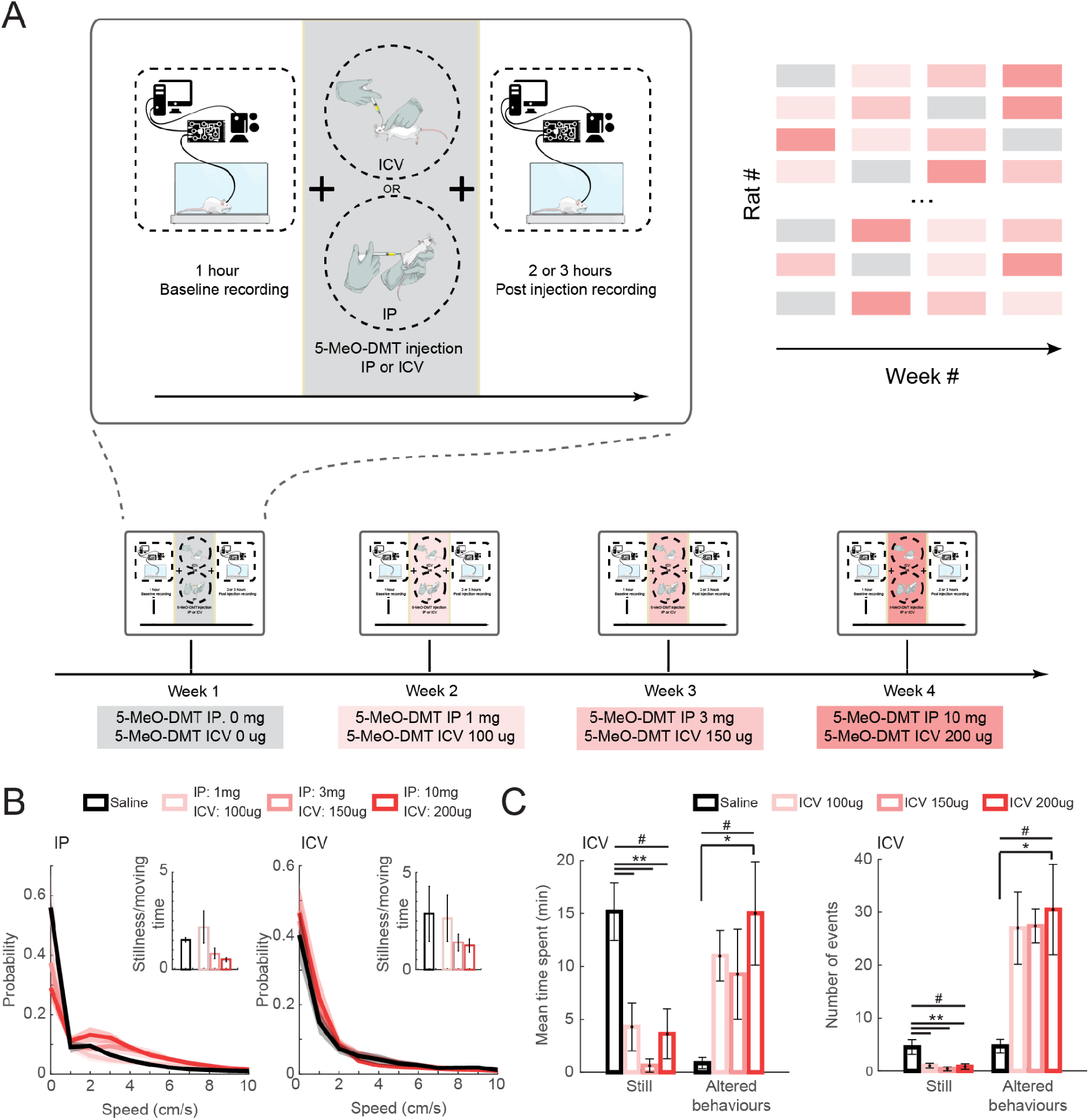
Experimental design and behavioural results. **(A)** Animals were submitted to 4 weeks of experiments for 5-MeO-DMT. On each experimental session we recorded 1h of baseline followed by 2 or 3 hours after drug injection; Two additional 1-hour recordings were made one day before and after the experiment but are not included in this work. Each week we treated animals with a different dose or vehicle (saline), in a randomised order among animals. **(B)** Normalised histogram of animal speed after IP (left) and ICV (right) injection of saline or different doses of 5-MeO-DMT. Insets show the average ratio between stillness (<1cm/s) and moving (between 1-8 cm/s). Error bars and shaded areas denote SEM. **(C)** Mean time spent (left panel) and the number of events (right panel) for the ‘Still’ behaviour and ‘Altered behaviours’ (see Supp. Figure 1). ANOVA #p<0.05, (‘Still’: Mean time: F(3,20)=7.6747, Number of events: F(3,20)=5.3164. ‘Altered behaviours’: Mean time spent: F(3,20)= 3.2362, Number of events: F(3,20)= 3.8690), with post hoc test *p<0.05; **p<0.01.

Before the first experimental session, animals were handled (∼20 min) and habituated to the experimental room and box for at least three days (1h per day). Experimental room temperature of 24°C was constant for all sessions. Experiments were done in the afternoon or beginning of the evening (14 h to 18 h) under dim light. For intracerebroventricular dosing, we used a Hamilton syringe of 10 ml connected to a polyethylene tube and a thinner cannula to fit in the implanted cannula. The volume dosing of 1ul unilaterally was manually administered for a duration of 30 s approximately.

### Recordings

The electrophysiological and video recordings were conducted using the Plexon Omniplex System or Intan Technologies coupled to a Logitech C920 webcam (30 Hz). Plexon and Intan signals were recorded at 40 kHz and 30 kHz, respectively, and downsampled to 1kHz for LFPs. Intan recordings were synchronised to video using a TTL pulse in a pseudorandom frequency to the Intan and a LED was placed in the filming area of the camera. No video alignment pre-processing was done for Plexon data since it is automatically synchronised. Animals were tracked using Cineplex from Plexon or IdTracker to detect the animals’ centre of mass. Once the tracking was performed, the animal’s speed was calculated. Behaviour analyses were done using Cineplex Editor.

### Power spectrum density estimates

We analysed the time-varying frequency content of the signal using a spectrogram. The signal was divided into 1-second overlapping segments, with a 50% overlap. The power spectral density (PSD) for each segment was calculated using Welch’s method, applying a Hamming window. The analysis used a sampling frequency of 1000 Hz and 4096 FFT points.

### Speed matching analysis

To address any potential behavioural differences between conditions, we employed a bootstrap procedure (Lopes-dos-Santos et al., 2018) to match the speed distributions of animals between conditions (baseline and post-injection sessions). Specifically, we randomly sampled half of the windows in a 30 min period post-injection and paired them with baseline windows based on the speed of the animals, thereby ensuring equivalent speed distributions across sessions. This procedure was repeated 500 times for each recording day. We then compared the mean delta (0.5-4 Hz) and theta (5-12 Hz) power of baseline and post-injection (eg. **Figure 2C, 2F, 2I, 2L**) conditions.

**Figure 2.**
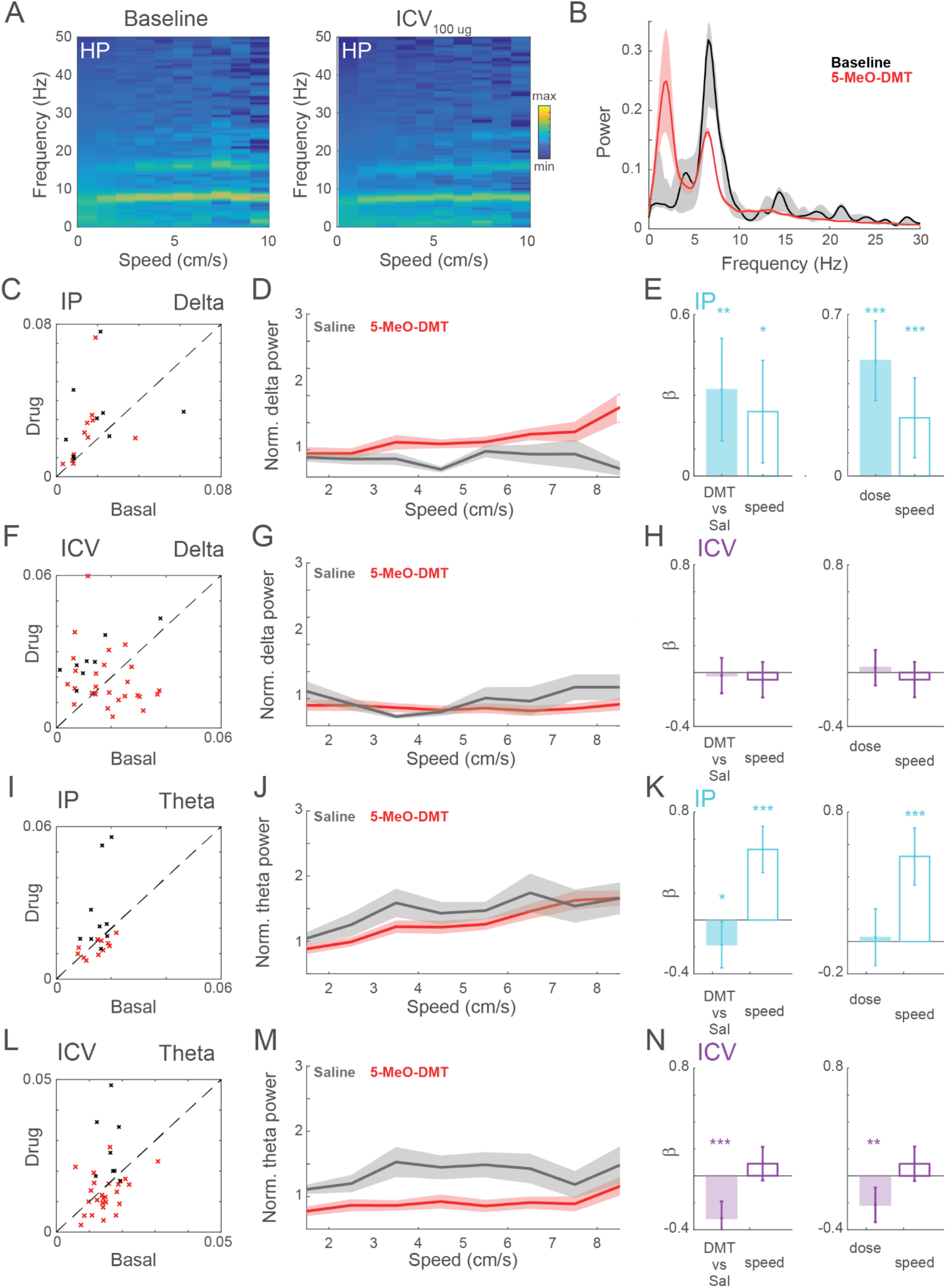
Changes in hippocampal delta and theta power after 5-MeO-DMT dosing. **(A)** Mean power spectrum at different bins of speed for a 30 min block of baseline (left) and 5-MeO-DMT at 100ug (right). **(B)** PSDs of one session obtained using the bootstrap procedure to equalise the speeds of drug and baseline periods (see methods). The shaded area represents the 95% confidence interval. **(C)** Mean delta power acquired from the bootstrap analysis during baseline and after IP dosing. Each cross corresponds to one experimental session after dosing (5-MeO-DMT: red; Saline: grey). **(D)** Mean normalised delta power of saline and drug conditions for each speed bin. **(E)** Coefficients of GML models estimated using the normalised power in D accounting for (left) animal speed and drug (DMT or saline), and (right) animal speed and normalised dose (0%: saline; 100%: maximum dose). **(F-H)** As in C-E, but for ICV injections. **(I)** Mean theta power acquired from the bootstrap analysis during baseline and after IP dosing. Each cross corresponds to one experimental session during the first 30 min after dosing (DMT: red; Saline: grey). **(J)** Mean normalised theta power of saline and drug conditions for each speed bin. **(K)** Coefficients of GML models estimated using the normalised power in D accounting for (left) animal speed and drug (DMT or saline), and (right) animal speed and normalised dose (0%: saline; 100%: maximum dose). **(L-N)** Similar to I-K, for ICV injections.

### GLM for assessing drug effect on delta and theta

To further investigate the effects of the drug on the power of delta and theta, we used a generalised linear model (GLM) approach. For each recording day, we computed the mean delta or theta power for different speed bins (1 cm/s wide bins with minimum values ranging from 1 to 8 cm/s). These power values were then divided by the mean obtained for the first speed bin (1-2 cm/s) in their corresponding baseline session. We used a linear regression model to predict the power values based on both the speed values and a categorical variable indicating whether the data were obtained from a 5-MeO-DMT or saline session. This approach allowed us to assess if the injection of 5-MeO-DMT was significantly different from saline administration after accounting for differences in speed. To assess for a potential dose-response relationship, we modified the model by replacing the binary categorical variable (5-MeO-DMT or saline) with the actual dose of the drug (with saline assigned a dose of 0). In all those analyses the channel with higher theta/delta ratio was used for both brain regions. The same analysis was done individually for both IP and ICV administrations.

### GLM for assessing drug effect on gamma oscillations

To investigate theta-nested gamma oscillations, we ran a matching analysis to determine whether any effects on gamma oscillations were in addition to what would be expected from theta power reductions and speed changes. To achieve this, we randomly selected theta cycles from the post-injection period and paired them with cycles from the baseline period (without repetition) that had similar speed and theta power. We defined a pair of cycles as “similar” when the difference between their speed and theta power values was less than 5% of the difference between the 10th and 90th percentiles of their overall distribution.

After matching the speed and theta distributions, we calculated the average gamma power of the paired theta cycles for each session. Subsequently, we normalised the average gamma power value obtained from the post-injection session by the mean value obtained from the baseline session (difference between the means divided by their sum). Finally, we fit a GLM as before, to predict relative gamma power from the speed and from categorical variables indicating whether each observation was obtained from a 5-MeO-DMT or saline session. Notice that this essentially is equivalent to a t-test but after matching the potential confounding variables (speed and theta power). Further, the same analysis was run but replacing this categorical variable by a specific dose (with saline assigned a dose of 0) to test for a dose-response relationship. These analyses were done individually for both IP and ICV administrations, and for slow and mid gamma oscillations.

To investigate the coupling between gamma amplitude and theta phase, we calculated theta-peak triggered averages of the (slow or mid) gamma envelope. Inconsistent relationships between gamma power and theta phase would result in flat averages, while consistent relationships would produce prominent theta components in these averages. Thus, quantify the strength of the coupling by calculating the power within the theta range (5-12 Hz) of these theta-peak triggered averages. The theta cycles used for this analysis were matched for speed and theta power, as in the previous analysis. To assess the effects of 5-MeO-DMT on this modulation, we used similar GLMs as the ones used to assess the effect on gamma power.

### Filtering theta and gamma oscillations

To extract the theta component, we filtered the LFP signals using a FIR filter for the band 5 - 12 Hz. To define the order of the filter we used the eegfilt function of the EEGLAB toolbox (Delorme and Makeig, 2004). Theta peaks were detected as local maxima of the filtered signals. To extract the envelopes of gamma oscillations, we first applied a similar FIR filter to each LFP signal with the corresponding gamma frequency band (slow: 25-50 Hz, mid: 50-100 Hz). We then estimated the envelope of each gamma by applying the Hilbert transform to the filtered signal.

### Awake-sleep cycle state maps

To study the wake and sleep states in the baseline, we used the spectral state map analysis described in Gervasoni et al., 2004 combined with the animal tracking obtained from the video recordings and data-driven thresholds of electrophysiological markers. Briefly, we first computed a spectrogram for each channel using a 1-s-window with 0.5 s of overlap. Then, we calculated two frequency ratios corresponding to delta and theta relative power; and gamma relative power (F <4.5Hz over F <=9Hz; and F< 20Hz over F<55 Hz). For each ratio, the first principal component was computed (across channels) and then used to plot the state map for the baseline of each recording session. From the video tracking (obtained from Cineplex or IdTracker, see above), we computed the mean animal velocity in the same timescale and periods of movement (>1 cm/s) were then used as landmarks to the identification of the wake stage in the state map. The region of each state (WK, SWS and REM) on the state map was then manually identified. For that, we also used the rest of the session to account for the possibility of not having sleep during the baseline, but always having the baseline period as a reference. Notice that the ‘canonical’ sleep staging can only be done for baseline periods, since the drug conditions may induce an altered state *per se*, that may not fit in the same classifications. The classification of REM-, SWS- and WK-like stages after the drug dosing was performed to characterise these altered states. However, it could be behaviourally attested that animals were, in general, moving after the first 15 min of drug dosing (i.e., were in a WK state).

Finally, we computed the mean duration of each wake-sleep state for saline and drug conditions, as well as the mean velocity of each state (across pooled episodes in **Figure 6 and Sup Figure 6**). We also computed the transition probabilities from going from each of the states to the others. For that analysis, unclassified periods with less than 20 s occurring between the same state were included in that state.

## RESULTS

### Behavioural effects of 5-MeO-DMT

First, we quantified the behaviours after 5-MeO-DMT ICV dosing (**Figure 1B, C; Supp Figure S1**). Besides some stereotyped behaviours already mentioned in the literature (e.g., *wetdog* shake), we quantified abnormal behaviours through empirical observation. More specifically, we show here the 10 behaviours most observed across animals: intermittent and uncoordinated gaiting or jumping; backwards gaiting; flat body gaiting; turning on its axis; quiet; uncoordinated gaiting; still; jumps; head tremor; and *wetdog* shake (**Supp Figure S1** and **Supplementary Table S1** for a more detailed description).

For each of those behaviours we measured the average time spent and the number of events (**Supp Figure S1**) during 5-MeO-DMT or saline ICV injection. We found that 5-MeO-DMT ICV sessions (all doses together) led to a significant increase of mean time spent and the number of events of the behaviour ‘intermittent uncoordinated gaiting or jumping’, ‘uncoordinated gaiting’ and ‘still’ (**Supp Figure S1**). To better understand and visualise the results we separated the behaviours in two groups: ‘still’ and ‘altered behaviours’ (a combination of behaviours 1 to 6), and evaluated the contribution of each dose to those two classes of behaviours (**Figure 1C**, ANOVA with post hoc test). We found that 5-MeO-DMT *vs* saline is significantly different for both behaviours (ANOVA, #p<0.05, ‘Still’: Mean time: F(3,20)=7.6747, Number of events: F(3,20)=5.3164. ‘Altered behaviours’: Mean time spent: F(3,20)= 3.2362, Number of events: F(3,20)= 3.8690). More specifically, both the mean time spent and the number of events of ‘still’ are significantly decreased for every dose compared to saline. Conversely, there was an increase of the mean time and number of events of ‘altered behaviours’ for the dose ICV 200 ug compared to saline (**Figure 1C**, Tukey-Kramer post hoc test, *p<0.05; **p<0.01).

#### Electrophysiological effects of 5-MeO-DMT

We next evaluated the impact of 5-MeO-DMT on the spectral profile of hippocampal LFPs. As the drug significantly affected locomotion (**Figure 1B**) and speed is known to influence LFPs, we employed a generalised linear model (GLM) to assess the effect of 5-MeO-DMT administration on delta (1-4 Hz) and theta (5-12 Hz) power while accounting for speed (**Figure 2**, see Methods for details). To achieve this, we first used the GLM to predict the relative delta or theta power (compared to the baseline session) while controlling for speed. Our analysis showed that there was a significant and dose-dependent impact of 5-MeO-DMT on delta power in the hippocampus following IP injections (**Figure 2C-E**, 5-MeO-DMT vs. saline: p=0.001; Dose: p<0.001). Interestingly, we did not observe the same effect for ICV injections. In contrast, we observed a significant decrease of theta power following the administration of 5-MeO-DMT using both IP (**Figure 2I-K**) and ICV (**Figure 2L-N**) delivery, with a significant dose-dependent effect specifically for ICV injections (DMT vs. saline: p<0.001; Dose: p=0.001). In summary, our findings indicate that 5-MeO DMT administration leads to a decrease in theta power in the hippocampus for equivalent locomotion speeds, as well as an increase in delta power specifically following IP injections.

Hippocampal theta oscillations are known to nest different gamma oscillations that reflect distinct circuits and pathways (for a review see (Fernandez-Ruiz et al., 2023). In alignment with previous studies (Belluscio et al., 2012; Fernández-Ruiz et al., 2017; Lasztóczi and Klausberger, 2016; Lopes-dos-Santos et al., 2018; Scheffer-Teixeira and Tort, 2017; Schomburg et al., 2014), we divided the gamma range in two components: slow (20-50 Hz) and mid (50-100). To assess the impact of 5-MeO-DMT administration on these two gamma rhythms, we conducted a matching analysis that controlled for changes in speed and theta power (**Figure 3A,B**; see Methods). This analysis revealed a significant dose-response relationship between 5-MeO-DMT ICV injection and slow gamma power (**Figure 3C**, DMT vs. saline: p=0.008; Dose: p=0.013). Specifically, slow gamma amplitude tended to be lower when nested in theta cycles during the post-injection epoch, compared to the baseline with matched power and speed. We observed a similar dose-response effect for mid gamma power (**Figure 3D**, DMT vs. saline: p=0.053; Dose: p=0.027). Notably, our results indicate that IP injections had no discernible effect on either slow or mid gamma power.

**Figure 3.**
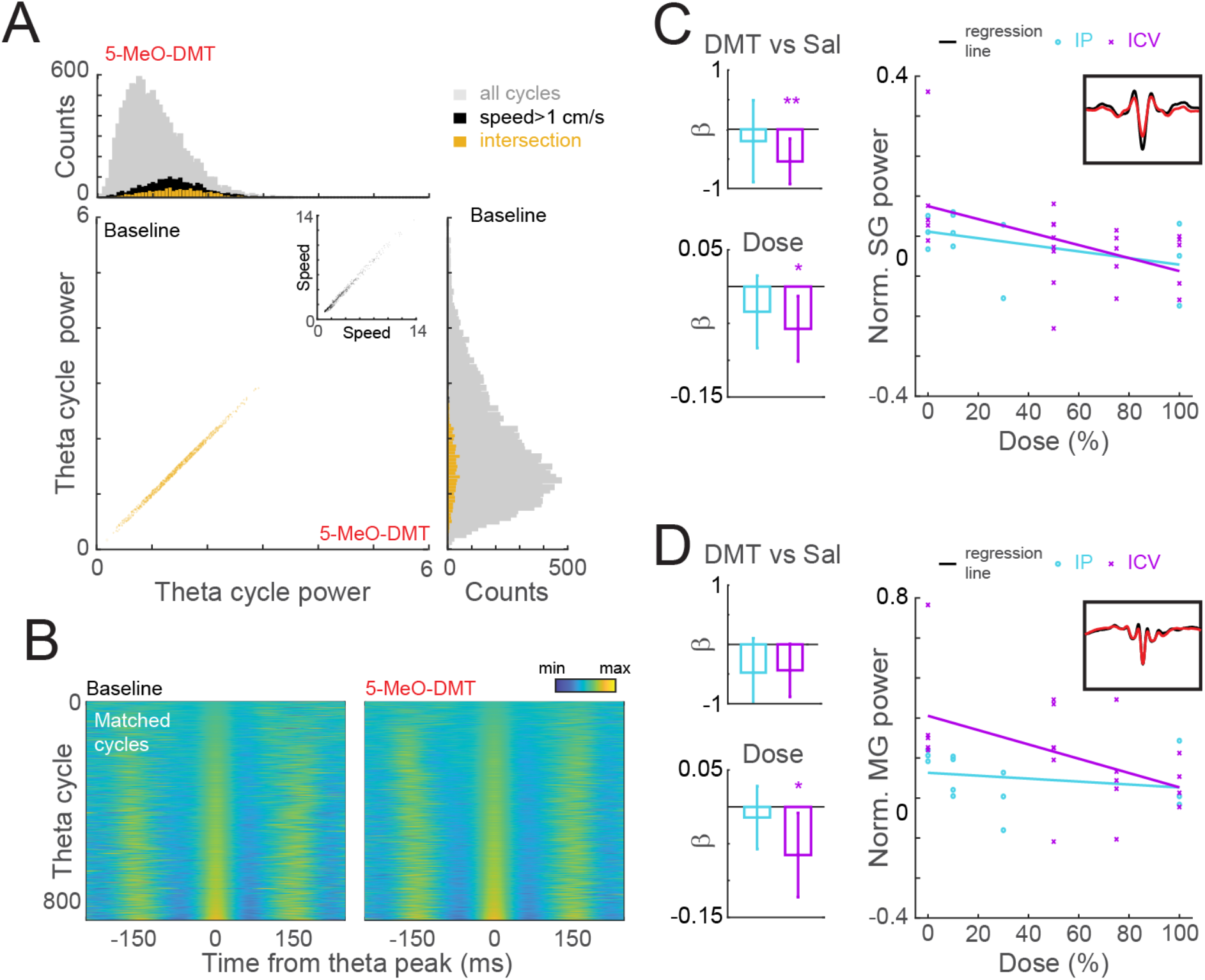
Slow- and mid-gamma power changes in the hippocampus after 5-Meo-DMT injection. **(A)** Example of theta cycle matching procedure. Each theta cycle occurring after the injection was matched to a theta cycle in baseline with a similar theta envelope and animal speed. **(B)** Representative recording day for matched theta cycles between baseline and post-injection conditions. Each row in the baseline panel corresponds to a distinct theta cycle, which is paired with a cycle of matched amplitude in the same row of the 5-MeO-DMT panel. The theta-filtered traces are depicted using pseudocolors and aligned to the corresponding theta peaks, providing a visual representation of the similarities between the two conditions. **(C)** The average slow-gamma envelope was then computed for each cycle in B. Bar plots show the coefficients of a GLM used to estimate slow-gamma power from the type of injection (DMT vs. saline; top) or from different doses of 5-MeO-DMT (bottom) as a variable. In the latter case, doses were normalised from 0 (saline) to 100% (maximal dose) (left). GLMs for IP and ICV were run separately. Errorbars denote 95% confidence interval and inset shows a representative slow-gamma triggered average. **(D)** Similar to C, but for mid-gamma power.

We proceeded to investigate the impact of 5-MeO-DMT on gamma amplitude modulation by the theta rhythm in the HP. To address the decrease in theta power caused by the drug we integrated theta amplitude into our matching analysis, enabling us to examine any additional effects on modulation beyond this reduction alone. Specifically, we selected individual theta cycles from pre- and post-injection epochs with matching amplitude and speed. We then calculated the average of the envelope of each gamma band, aligned to those theta peaks (see **Figure 4A,C**). To evaluate theta modulation, we computed the power spectra of the triggered average envelopes (see methods) and quantified the power within the theta range. A higher concentration of power within the theta range of the triggered envelopes indicates a stronger modulation between the theta phase and the gamma amplitude.

**Figure 4.**
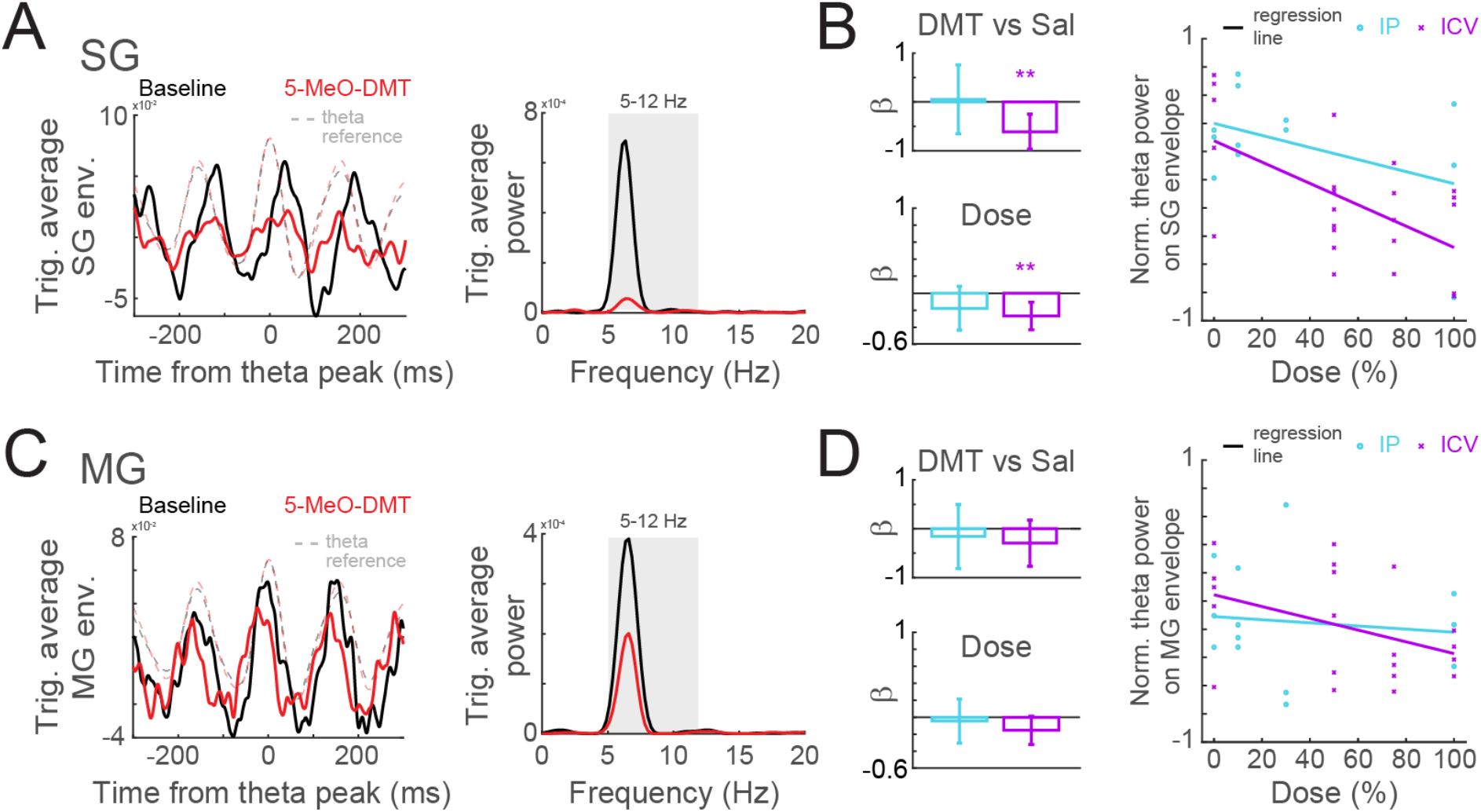
Hippocampal theta-nested slow- and mid-gamma oscillations after 5-MeO-DMT. **A.** (Left) Average of slow-gamma envelope, triggered by theta peaks (dashed lines) after 5-MeO-DMT injection (red) and during baseline (black). (Right) Power spectrum of the triggered average on the left. Modulation was estimated by the average power in theta frequency band (shaded area). Theta cycles from baseline were matched in both mean theta amplitude (average envelope) and animal speed to account for the influence of those variables into gamma coupling. **B.** Coefficients of GLM of theta-SG modulation, estimated using the injected drug (top) and taking the dose into account (bottom, right) for IP (light blue) and ICV (magenta) injections. GLMs for IP and ICV were computed separately. **C-D.** Same as A-B, but for mid-gamma.

Following this analysis, we found that 5-MeO-DMT administration resulted in a significant reduction in the theta modulation of slow gamma oscillations specifically after ICV administration (DMT vs. saline: p=0.002; Dose: p=0.003). Furthermore, we found no effect on the theta modulation of mid gamma oscillations in the HP.

Next, we investigated the impact of 5-MeO-DMT on prefrontal cortex delta and theta power (**Figure 5**) using the same analysis framework for the HP channels (**Figure 2**). Our findings indicate that IP injections of 5-MeO-DMT led to a dose-dependent increase in delta (DMT vs. saline: p<0.001; Dose: p<0.001) and theta power (DMT vs. saline: p=0.002; Dose: p<0.001), while ICV administration resulted in a significant dose-dependent decrease of theta power (DMT vs. saline: p<0.001; Dose: p<0.001). These opposite effects suggest that the route of administration plays a critical role in the impact of 5-MeO-DMT on prefrontal cortex delta power. Further, we observed no significant effect for the theta range with IP injection, whereas ICV administration led to a dose-dependent decrease of theta power, consistent with the HP results.

**Figure 5:**
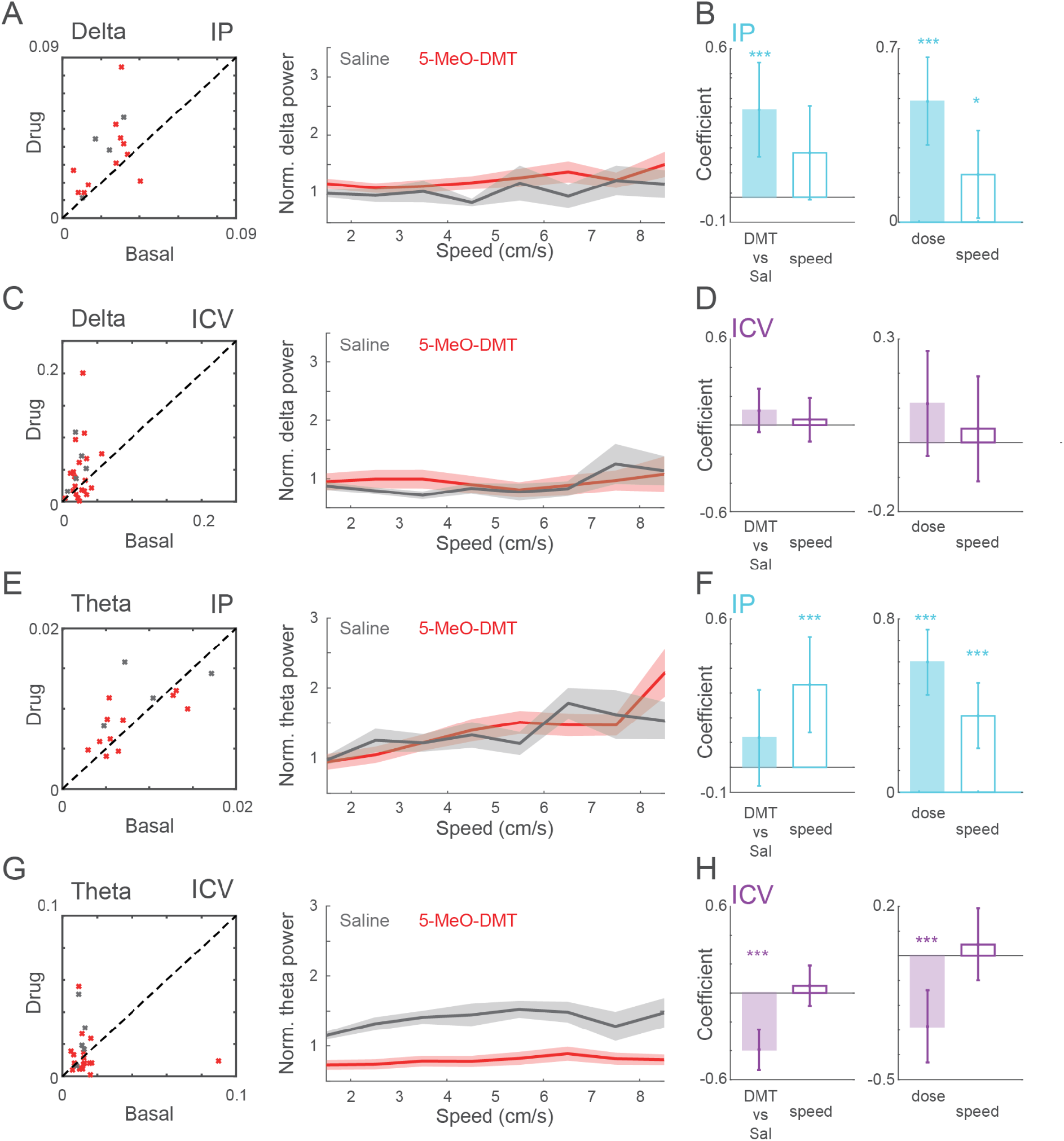
Changes in mPFC spectral power after IP and ICV 5-MeO-DMT dosing. **(A)** (Left) Mean delta power acquired from the bootstrap analysis during baseline and after dosing. Each cross corresponds to one experimental session (DMT: red; Saline: grey) after dosing. (Right) Mean normalised delta power of saline and drug conditions. **(B)** GLM delta power for IP dosing. **(C and D)** Similar to A and B, for theta band. **(E-H)** Similar to A-D, for ICV dosing. Notice the decrease in theta normalised power after 5-MeO-DMT in comparison to saline dosing.

To analyse the spectral patterns of 1-second windows of combined hippocampal and cortical activity, we used a manifold approach called *state map* (Gervasoni et al., 2004). This approach generates a two-dimensional representation of LFPs by calculating the ratio of two frequency bands: (1) delta over theta plus delta, and (2) from delta to beta over delta to gamma (see Methods for details). The state map method was primarily designed to identify three main clusters of the sleep-wake cycle: awake, SWS, and REM. In summary, high amplitude theta power (ie, low ratio 1) distinguish awake and REM from SWS, whereas relative gamma power on theta oscillations separates awake from REM (awake theta cycles exhibit higher gamma power compared to REM cycles, captured by ratio 2).

We first used the state maps of the baseline session of each animal to define the three main sleep-awake clusters (see **Supplementary Figure S3-4** for examples). Then, we projected the activity of post-injection windows onto that map (**Figure 6A** and **Supp Figure S2A**). Consistent with our previous results, this analysis revealed that animals exhibited an altered spectral profile after 5-MeO-DMT dosing. Once we had the state map for pre- and post-dosing periods for saline and drug experiments, we calculated the transition between each pair of states (**Figures 6B**). We found that saline and 5-MeO-DMT groups had similar transition probabilities (**Figures 6C** for ICV, **Supp Figure S2B** for IP). Notwithstanding, transitions between WK-like and REM-like states tended to increase for 5-MeO-DMT ICV experiments - especially for the doses 100ug and 150ug (**Figure 6B**; see the increase in the mean of transition probability for the WK purple bars in **Figure 6C**).

**Figure 6:**
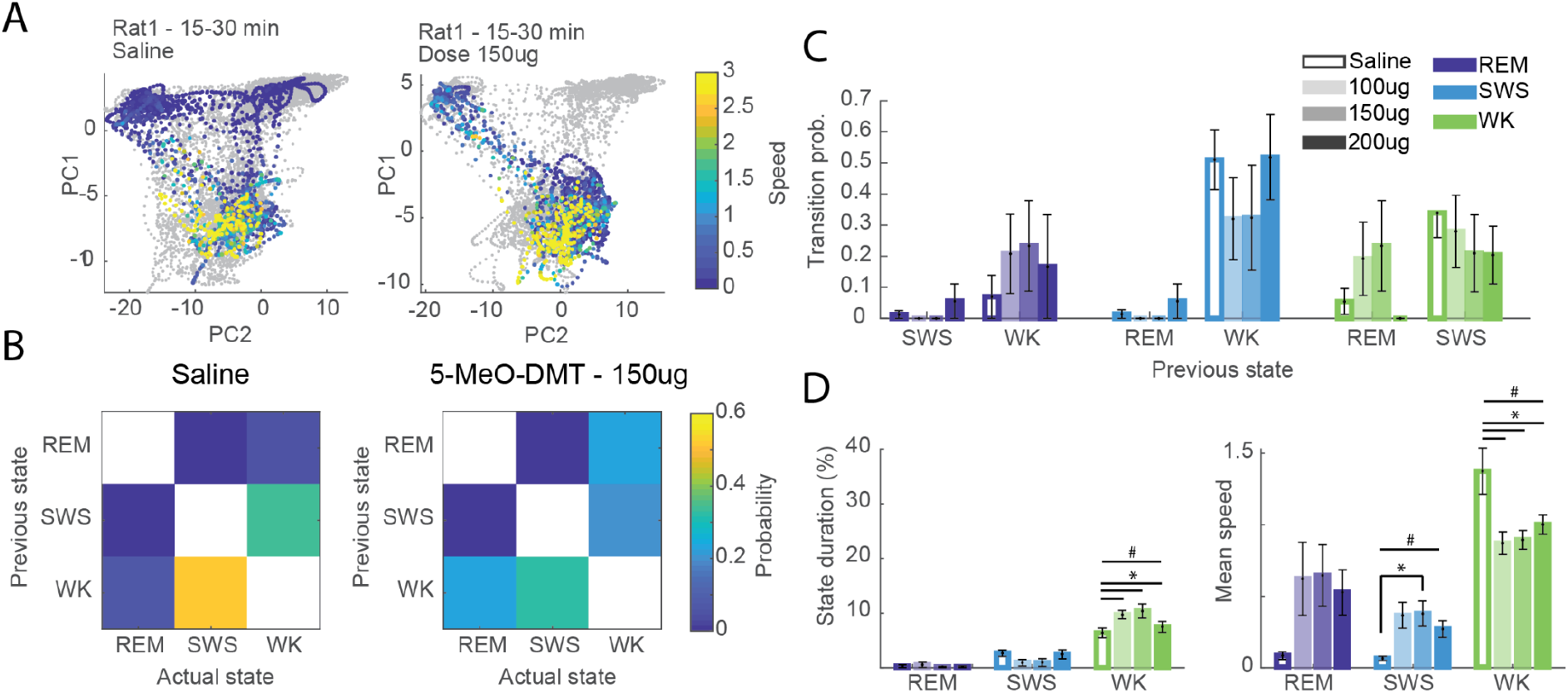
State map spectral changes resemble the sleep-wake cycle after ICV 5-MeO-DMT dosing. **(A).** Representative examples of state maps of 5-MeO-DMT experiments. Grey trace represents the baseline period. **(B)** Transition matrix showing the probability of change between each pair of states for saline (left panel) and 5-MeO-DMT (from left to the right panel: 100ug, 150ug and 200ug) experiments. **(C)** Transition probability to REM (dark blue), SWS (light blue) and WK (green), given the specified previous state (x-axis). Empty bars for saline and filled bars for 5-MeO-DMT at doses 100ug, 150ug and 200ug (from lighter to darker colours). **D.** Percentage of state duration (left panel) and mean speed (right panel) of each sleep-waking state for saline and different doses of 5-MeO-DMT experiments (ANOVA, #p<0.05 (WK: State duration: F(3,27)=3,638); SWS: Mean speed: F(3,320)=3,0864; WK: Mean speed: F(3,1560)=4,7072), post hoc test *p<0.05, SEM; n= episodes of sleep or wake).

Next, we computed the duration and animal speed (locomotion) for each sleep or wake episode. We found a significant increase in the time spent in WK state after 5-MeO-DMT ICV dosing, while it did not change for REM-like or SWS-like states compared to saline (**Figure 6D**, left panel). Conversely, we found that mean speed across episodes was increased during REM-like and SWS-like states, while it decreased during WK after 5-MeO-DMT IP (**Figure 6D**, right panel). Animals did not spend more time in a sleep state; they rather presented a greater speed during that sleep-like state. In that sense, we could say that there was a significant change in the state map profile that was not accompanied by the behavioural state. In other words, 5-MeO-DMT administration pushed the hippocampal activity closer to the states observed during baseline sleep, as indicated by post-5-MeO-DMT periods invading the SWS and REM clusters in the state map. Note in the representative examples the abnormal transitions (**Figure 6B**, left panel) and high speed cluster displacement from WK, an effect not seen for saline (**Figure 6D** right).

We found similar state map changes for the 5-MeO-DMT IP dosing, even though the transition probability between sleep-like and wake-like clusters did not change significantly (**Supp Figure S2B**), Likewise in ICV experiments, in the IP experiments there was an increase in time spent in WK and a decrease in the WK mean speed across episodes (**Supp Figure S2C**). The opposite occurred for the SWS-like state, for animals spent more time and at higher speed within the SWS-like state after 5-MeO-DMT IP dosing (**Supp Figure S2C**). In other words, the rats presented a SWS-like state even though they were actively moving across the open field. A similar yet non-significant trend occurred during the REM-like state; the REM-like regions at the dose of 10 mg had higher speed compared to saline (**Supp Figure S2C**, right panel). In this case, animals not just presented a displacement of higher speeds cluster, but also such displacement occurred during a longer period and it seems to be towards the SWS-like cluster (or even to REM-like). Note that effect in the examples shown in **Supp Figure S2A**. Rats 3-5 presented a high-speed cluster closer to SWS-like compared to Rat5 during saline (**Supp Figure S2A**, right-bottom panel).

## DISCUSSION

This study provided the first behavioural and electrophysiological quantification of speed-controlled LFP simultaneously recorded from the HP and the mPFC of freely behaving rats subjected to an acute psychedelic experience induced by 5-MeO-DMT at various doses.

### Behavioural changes induced by 5-MeO-DMT

Animals treated with 5-MeO-DMT exhibited a variety of altered behaviours (**Figure 1, Supp Fig S1**), some of which are well documented stereotyped behaviours, while others were not previously reported in the literature. This may be attributed to the fact that most studies assess behavioural effects of drugs on specific tasks, rather than on spontaneous activity. Our findings indicate that the most significant changes in behaviour were related to ‘uncoordinated gaiting’ and ‘stillness’.

The ‘Uncoordinated gaiting’ behaviour observed in our study may be related to the ‘forepaw treating’ associated with serotonergic syndrome (Haberzettl et al., 2013). Spanos et al. showed a dose-dependent increase in “forepaw treating” and ‘low body posture’, but not in ‘head weaving’ after MDMA injection in rats (Spanos and Yamamoto, 1989). Conversely, we did not observe a significant change in the behaviour equivalent to ‘flat body gaiting’ or ‘low body posture’.

In accordance with the appearance of altered behaviours, we found that animals exhibited a lower level of stillness (both in duration and frequency) following administration of 5-MeO-DMT at all doses compared to when they were given saline (**Figure 1B and C**).

Although the remaining quantified behaviours did not change individually, they did show a general increase in prevalence following administration of 5-MeO-DMT. When grouping behaviours 1 to 6 as “altered behaviours,” we observed a significant increase in both the duration and frequency of these behaviours (**Figure 1C**). This effect appeared to be dose-dependent, with a stronger impact observed at higher doses of the drug. While the behaviours of ‘backward gaiting’, ‘head tremor’ and *‘wetdog* shake’ did not show significant changes when considered individually, we observed an increase in their prevalence when analysed together, consistent with observations reported in mice treated with d-LSD (Kyzar et al., 2016). *Wetdog* shake, also known as head-twitch response (HTR), is a rapid and sudden side-to-side shaking of the head/neck that is a well-known marker of the agonistic effect of the serotonergic system (Bedard and Pycock, 1977; Halberstadt and Geyer, 2013). This behaviour is highly rhythmic and produces a wave-like oscillation around 90 Hz in mice (Halberstadt and Geyer, 2013). In this study, we observed the occurrence of ‘wetdog shake’ in rats treated with 5-MeO-DMT (**Supp Figure S1B**), which is consistent with evidence indicating that HTR is increased by other psychedelics in mice, such as DOI, DOM, and N,N-DMT (Halberstadt et al., 2020). However, we did not observe a significant change in the flat body gait behaviour (**Supp Figure S1**), as previously reported for d-LSD (Kyzar et al., 2016). Likewise, we found no significant increase in the duration of ‘turning around its axis’, a behaviour that has not been well described in the current literature on psychedelics, although 5-MeO-DMT tended to increase the number of events.

The differences observed between ICV and IP dosing may be related to the availability of 5-HT agonists, peripheral action, and the amount of drug that reaches the central nervous system. Our IP findings contrast with a similar study by Halberstadt et al., 2008, which found no initial increase in locomotion, but reported a later hyperactivity (Halberstadt et al., 2012, 2008). However, it is important to note that lower doses were used in their study, potentially leading to a subliminal or delayed effects on locomotion. The increase in locomotion we observed in the present study could be an effect of higher doses and more direct route of administration in the ICV experiments.

### LFP power changes induced by 5-MeO-DMT

We found a significant decrease of hippocampal theta power in a dose-response manner that could not be explained by changes in animal speed. This is in accordance with the idea that the median raphe acts as a theta desynchronization nucleus (Jackson et al., 2008; Vertes, 1981; Vinogradova et al., 1999). Moreover, the power of theta-associated gamma oscillations and their coupling to theta phase were significantly reduced by 5-MeO-DMT. The administration of 5-MeO-DMT induced significant alterations in animal locomotion, which is an important factor to consider when interpreting the potential effects observed in electrophysiology, as speed can have a profound impact. It is worth noting that our study differs from previous investigations in that we took this confounding variable into consideration.

### Differences between IP versus ICV experiments

The differences between IP and ICV dosing are also noteworthy. In the hippocampus, the dose-dependent increase in delta power following IP injections was not observed for ICV injections, implying a possible peripheral mechanism. Classical psychedelics such as 5-Meo-DMT can also bind to serotonergic receptors of the peripheral nervous system (PNS) and this could affect the CNS quite differently from the ICV dosing. Serotonin is a major controller of the gut motility, including intrinsic reflex, epithelial secretions, and vasodilatation. Furthermore, its signalling takes part in vagal extrinsic and spinal afferent fibre activation, leading to pancreatic secretion, satiation, pain and discomfort, nausea, and vomiting (Mawe and Hoffman, 2013). The high availability of serotonin in the PNS provided by the IP dosing may have led to a peripheral increase in receptor activation, causing physiological alterations such as gastric discomfort. Although not life-threatening, those alterations could potentially change the subjective experience of the animal during the experience, as well as its neural correlates. A second non-exclusive hypothesis is that the peripheral binding of psychedelics causes fewer molecules to reach the CNS. Yet, these putative lower doses were still effective, as shown by the significant electrophysiological changes observed in the HP and mPFC for theta and gamma oscillations after IP injections of 5-MeO-DMT. Regarding theta-nested gamma oscillations, IP injections produced less dose-dependency than ICV injections, which suggests that IP injections either dampen or shift the dynamic range of the effective intracerebral doses within the HP or mPFC.

### Changes in sleep and waking states induced by 5-MeO-DMT

State maps based on LFP power ratios provide a consistent measure of state-dependent spectral features as animals alternate between sleep and waking (Gervasoni et al., 2004). Under drug-free conditions, state maps usually display very similar patterns across animals, despite inter-individual differences in behaviour and LFP signals. In the present study the state maps presented marked alterations after psychedelic dosing compared to saline, with some changes shared by the two groups investigated (**Figure 6 and Supp Figure S2**). More specifically, ICV 5-MeO-DMT dosing did not influence the duration of SWS-like episodes but rather increased the animals’ speed during this spectral state (**Figure 6C**). This result indicates that the animals treated with ICV 5-MeO-DMT were behaviourally awake, but their HP seems to be in an SWS-like state. There was also an increase in the duration of WK episodes, although the animals’ speed decreased after ICV 5-MeO-DMT (**Figure 6C**). Therefore, when animals had electrophysiological features of WK, they were more likely to be at a lower speed after ICV 5-MeO-DMT compared to saline.

The results obtained in the IP 5-MeO-DMT experiments were overall like those from the ICV 5-MeO-DMT experiments, with animals spending more time in the SWS-like and WK-like spectral regions (**Supp Figure S2C**, left panel), and presenting higher speeds during the SWS-like state and lower speeds during the WK-like state, when compared to animals treated with saline (**Supp Figure S2C**, right panel).

Overall, the results presented here strengthen the notion that psychedelics induce significant changes in the sleep-wake cycle, possibly leading to sleep-like waking states of hippocampal LFP oscillations (Bréant et al., 2022). They also resonate with early studies of the effects of d-LSD in humans. Muzio et al., (1966) found that low doses (6-40 ug) caused a prolongation of the first and second REM sleep episodes, and a shortening of subsequent episodes, with brief REM sleep episodes interrupting SWS. Another study showed a decrease in latency between the bouts of REM sleep after dosing, with a decrease in theta activity (Torda, 1968). Altogether, the available evidence suggests that the psychedelic experience precludes sleep at the behavioural level yet allows the brain to continue cycling across waking-like and sleep-like electrophysiological states.

It is important however to consider that the LFP analysis of the WK-like, SWS-like and REM-like spectral regions of state maps does not comprise the behavioural parameters that characterise sleep-wake states, such as immobility or muscle atonia. Furthermore, the SWS-like or REM-like spectral regions are not equivalent to the SWS or REM spectral regions, but rather reflect proportional spectral features that point to similar frequency patterns across sleep states. Further studies shall clarify whether the sleep-like electrophysiological features induced by psychedelics are indeed related to the dream-like psychological features of the psychedelic experience.

## Funding

Work supported by CAPES and CNPq.

## Acknowledgments

We thank S Ruschi, F Cini, I Gendriz, DA Filho, D Golbert, C Pereira, V Cota, W Blanco, J Oliveira, V Andino, D de Araujo, S Rehen, LF Tofoli, D Laplagne for insightful discussions, K Rocha for administrative support, G Santana for secretarial assistance, AE Oliveira for miscellaneous help.

## Author contributions (names must be given as initials)

**Conceptualization & Experimental Design:** SR, VLS, ACS; **Data Collection:** ACS, VLS, AF, MM, NCS; **Data Analysis:** BCS, ACS, VLS; **Figure Conceptualization & Preparation:** ACS, BCS, VLS, SR; **Paper Writing:** ACS, BCS, SR, VLS; **Paper revision:** AF, MM, NCS, KL, ABLT, RL; **Project Supervision:** SR, VLS.

## Data availability statement

The data will be made available upon reasonable request.

## Additional Information (including a Competing Interests Statement)

The authors declare no competing interests.

## SUPPLEMENTARY MATERIAL

**Supplementary Table S1:** Description of the behavioural analysis for the first 30 min after 5-MeO-DMT (ICV) or saline dosing.

|  | Behaviour | Description |
| --- | --- | --- |
| 1 | Intermittent and uncoordinated gaiting or jumping | The animal displayed an intermittent behaviour of walking and stopping within short periods of interval (~10s). While walking its paws presented an uncoordinated rhythm, and/or it moved by making sudden jumps forward. |
| 2 | Backward gaiting | The animal walked backwards with short/small or long steps |
| 3 | Flat body gaiting | Animal displayed gaiting with lower posture, with his body (ventral core) collapsed to the floor |
| 4 | Turning on its axis | The animal turned circularly with its head toward its tail basis, or its forepaws treading were making it turn to the side without moving its back paws. |
| 5 | Quiet | Periods when the animal was quieter than active awake, but not completely still |
| 6 | Uncoordinated gaiting | Animal walking with its paws in an uncoordinated rhythm |
| 7 | Still | Animal completely still, very quiet with no movement. |
| 8 | Jumps | Sudden jumps forward |
| **9** | Head tremor | The animal presented a head-tremor different from the wetdog shake. In a slower rhythm, as if it was biting its teeth |
| **10** | Wetdog shake | The animal presented a head-twist from side to side very sudden, rapid with great lateral amplitude. |

**Supplementary Figure S1:**
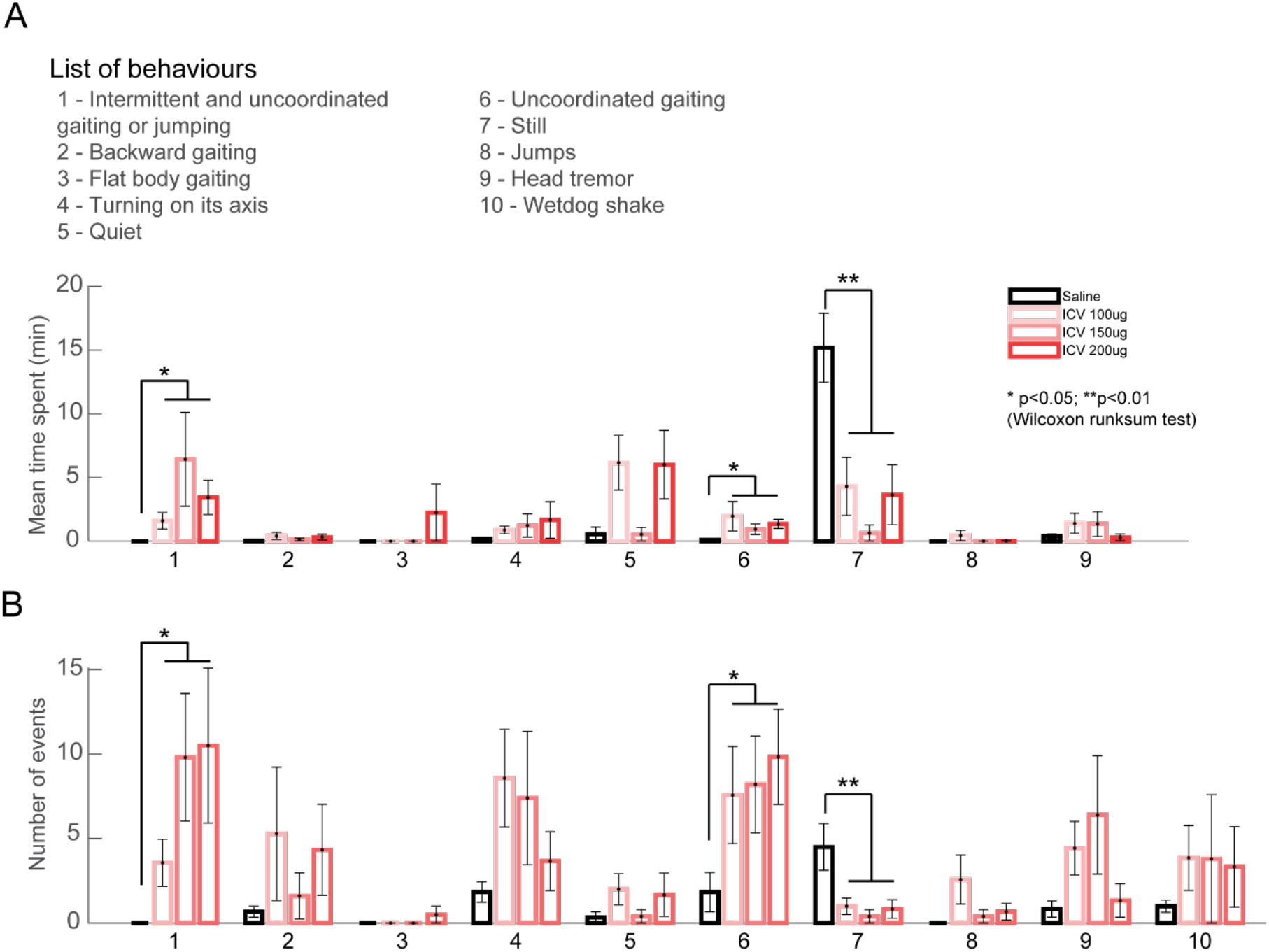
Behaviours are altered after ICV 5-MeO-DMT dosing. **(A)** Mean time spent in each behaviour (1 to 9) for ICV 5-MeO-DMT experiments for all ICV doses (100ug, 150ug and 200ug). **(B)** The number of events of each behaviour (1 to 10) for ICV 5-MeO-DMT experiments for all doses (100ug, 150ug and 200ug). *p<0.05, **p<0.01, Wilcoxon rank-sum test for all conditions drug *versus* saline.

**Supplementary Figure S2:**
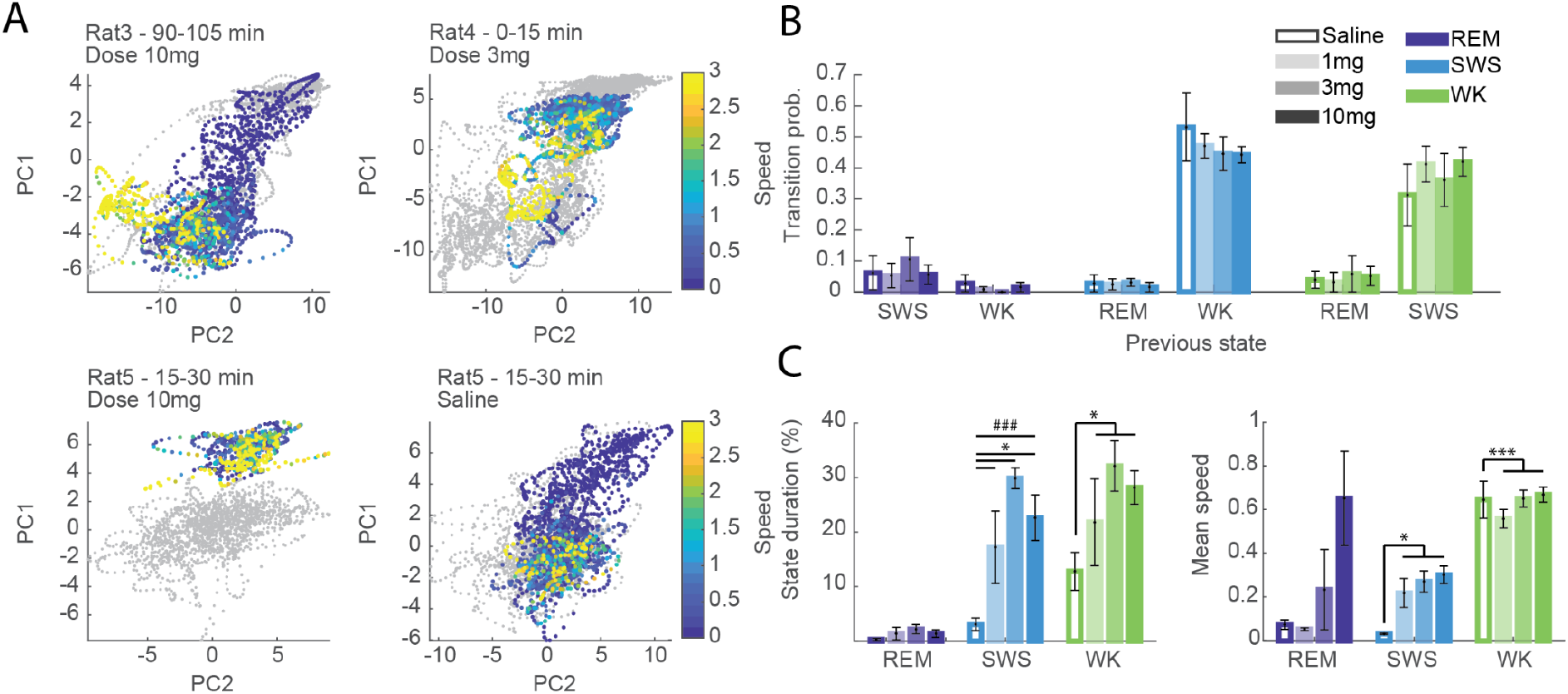
State map changes after IP 5-MeO-DMT dosing. **(A).** Representative examples of state maps of 5-MeO-DMT experiments. Grey trace represents the baseline period; black refers to the post-dosing period and red to post-dosing periods of animal’s speed >1 cm/s. Analyses performed for the post-dosing period of 30min (and until 3h30min for some animals). **(B)** Transition probability to REM (dark blue), SWS (light blue) and WK (green), given the specified previous state (x-axis). Empty bars for saline and filled bars for 5-MeO-DMT at doses 1mg, 3mg and 10mg (from lighter to darker colours). **C.** Percentage of state duration (left panel) and mean speed (right panel) of each sleep-waking state for saline and different doses of 5-MeO-DMT experiments (State duration: ANOVA, ###p<0.001 (SWS: State duration: F(3,13)=10,3419), post hoc test, *<0.05, SEM; Mean speed: Wilcoxon rank-sum test, *p<0.05 ***p<0.001, SEM; n= episode of sleep or wake).

**Supplementary Figure S3:**
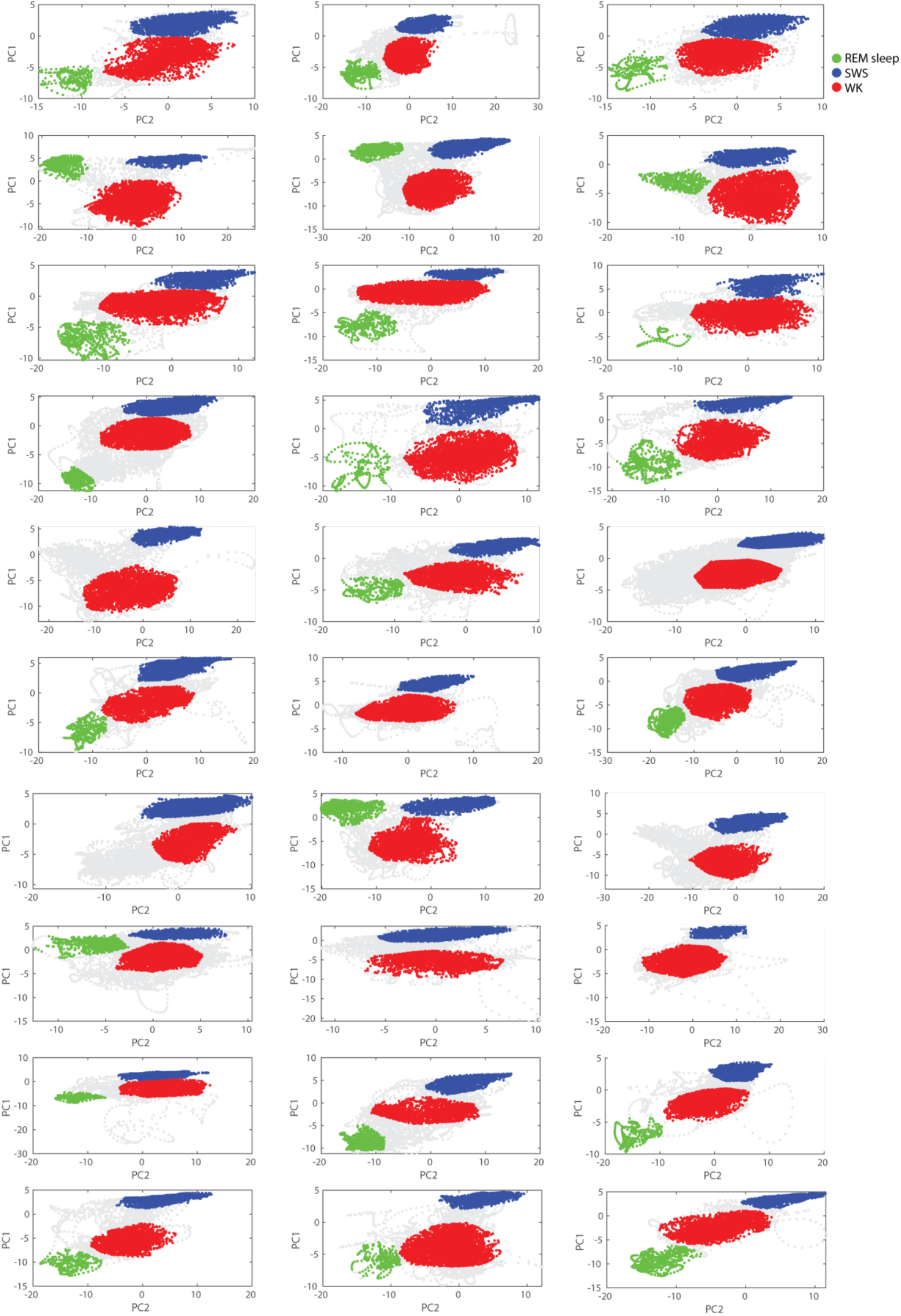
Manual annotations of sleep states from the 2D spectral maps for 5-MeO-DMT ICV experiments. Each map is delimited for the baseline session and then the experimental period is assigned based on the baseline map previously defined. The assignments of waking periods were guided by the occurrence of high animal’s speed. Both baseline and drug periods are shown in this figure. Each dot corresponds to 1 second period. WK, SWS and REM sleep periods are shown in red, blue, and green, respectively.

**Supplementary Figure S4:**
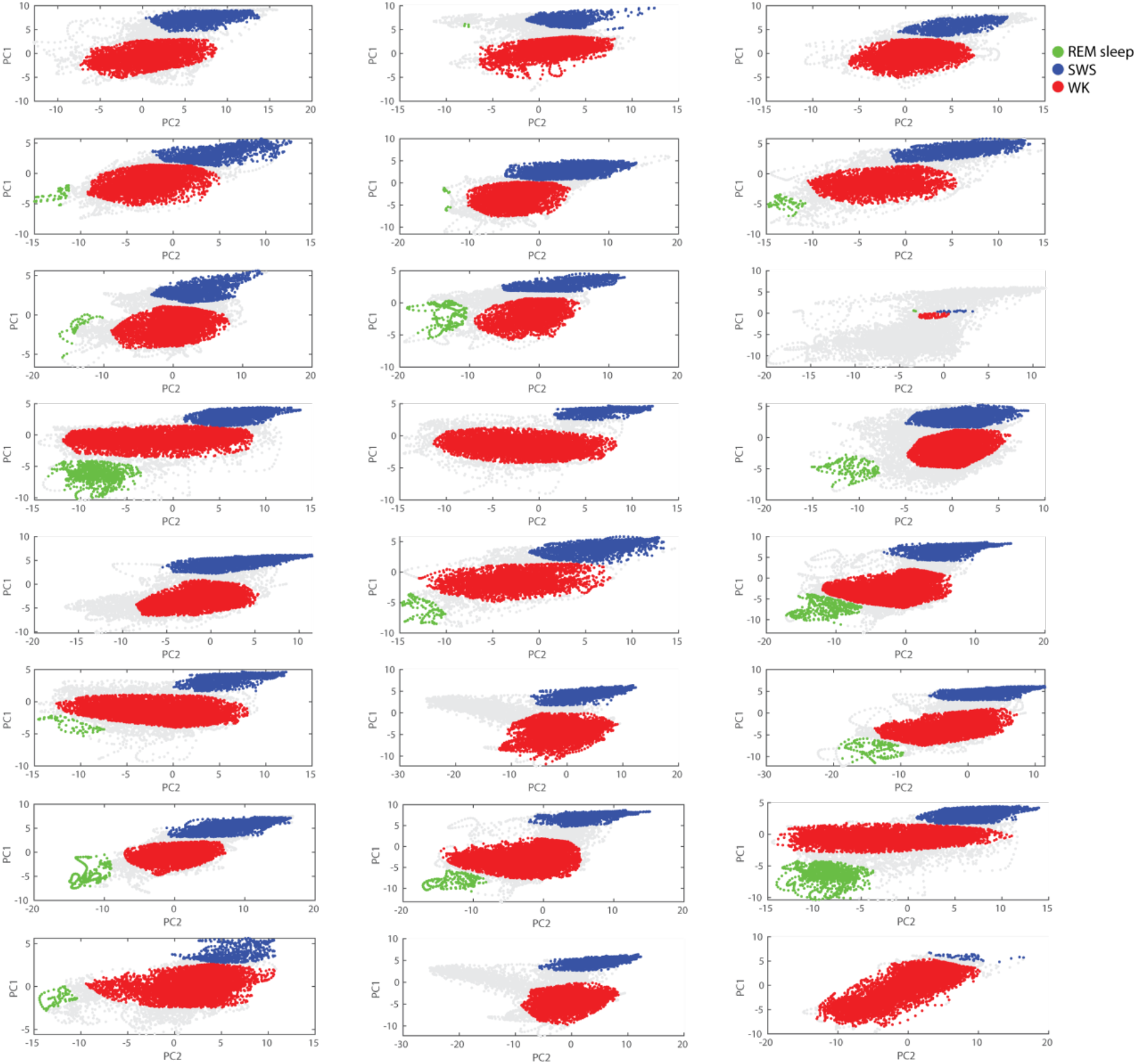
Manual annotations of sleep states from the 2D spectral maps for 5-MeO-DMT IP experiments. Each map is delimited for the baseline session and then the experimental period is assigned based on the baseline map previously defined. The assignments of waking periods were guided by the occurrence of high animal’s speed. Both baseline and drug periods are shown in this figure. Each dot corresponds to 1 s period. WK, SWS and REM sleep periods are shown in red, blue, and green, respectively.

## Notes

### Competing Interest Statement

The authors have declared no competing interest.

